# The mechanics of cephalic furrow formation in the *Drosophila* embryo

**DOI:** 10.1101/2023.01.19.524786

**Authors:** Redowan A. Niloy, Michael C. Holcomb, Jeffrey H. Thomas, Jerzy Blawzdziewicz

## Abstract

Cephalic furrow formation (CFF) is a major morphogenetic movement during gastrulation in *Drosophila melanogaster* embryos that gives rise to a deep, transitory epithelial invagination. Recent studies have identified the individual cell shape changes that drive the initiation and progression phases of CFF; however, the underlying mechanics of these changes are not yet well understood. During the progression phase, the furrow deepens as columnar cells from both the anterior and posterior directions fold inwards rotating by 90°. To analyze the mechanics of this process, we have developed an advanced 2D vertex model, which introduces multi-node representation of cellular membranes and allows us to capture the membrane curvature associated with pressure variation. Our investigations reveal some key mechanical features of CFF. As cells begin to roll over the cephalic furrow cleft, they become wedge-shaped as their apical cortices and overlying membranes expand, lateral cortices and overlying membranes release tension, internal pressures drop, and basal cortices and membranes contract. Cells then reverse the process by shortening apical cortices and membranes, increasing lateral tension, and causing internal pressures to rise. Since the basal membranes expand, the cells recover a rotated columnar shape at the end of this process. Interestingly, our findings indicate that the basal membranes may be passively reactive throughout the progression phase. We also find that the smooth rolling of cells over the cephalic furrow cleft necessitates that internalized cells provide a solid base through high membrane tensions and internal pressure levels, which allows transmission of tensile force that pulls new cells into the furrow. These results lead us to suggest that CFF may help establish a baseline tension across the apical surface of the embryo that would facilitate cellular coordination of other morphogenetic movements via mechanical stress feedback mechanisms.

**SIGNIFICANCE:** Mechanical forces and stress feedback are essential for the development of morphology and structure in the embryo. Although great progress has been made in understanding the genetic control of patterning and cell fate, mechanical stress contributions are not as well understood. Mechanical analyses of the apical constrictions initiating ventral furrow formation and subsequent invagination dynamics in *Drosophila* have shed considerable light on these processes; however, ventral furrow formation is only one of many morphogenetic movements. Cephalic furrow formation occurs simultaneously with ventral furrow formation, but its cell shape changes and invagination dynamics are radically different. This study shows that mechanical forces and feedback operating in cephalic furrow formation also differ considerably from those in ventral furrow, demonstrating a potentially wide array of mechanical processes in morphogenesis.

## INTRODUCTION

Gastrulation in the *Drosophila* embryo involves several major invaginations, most pronounced of which are the ventral furrow (VF), posterior midgut (PMG), and cephalic furrow (CF) invaginations (1, 2). CF begins to grow just after the initiation of VF formation (VFF). The CF cleft first appears on the lateral sides of the embryo at around 65% of the egg length; as CF develops, it extends along the entire embryo perimeter (Fig. 1 *a* and 1 *b*), (3, 4).

**Figure 1:**
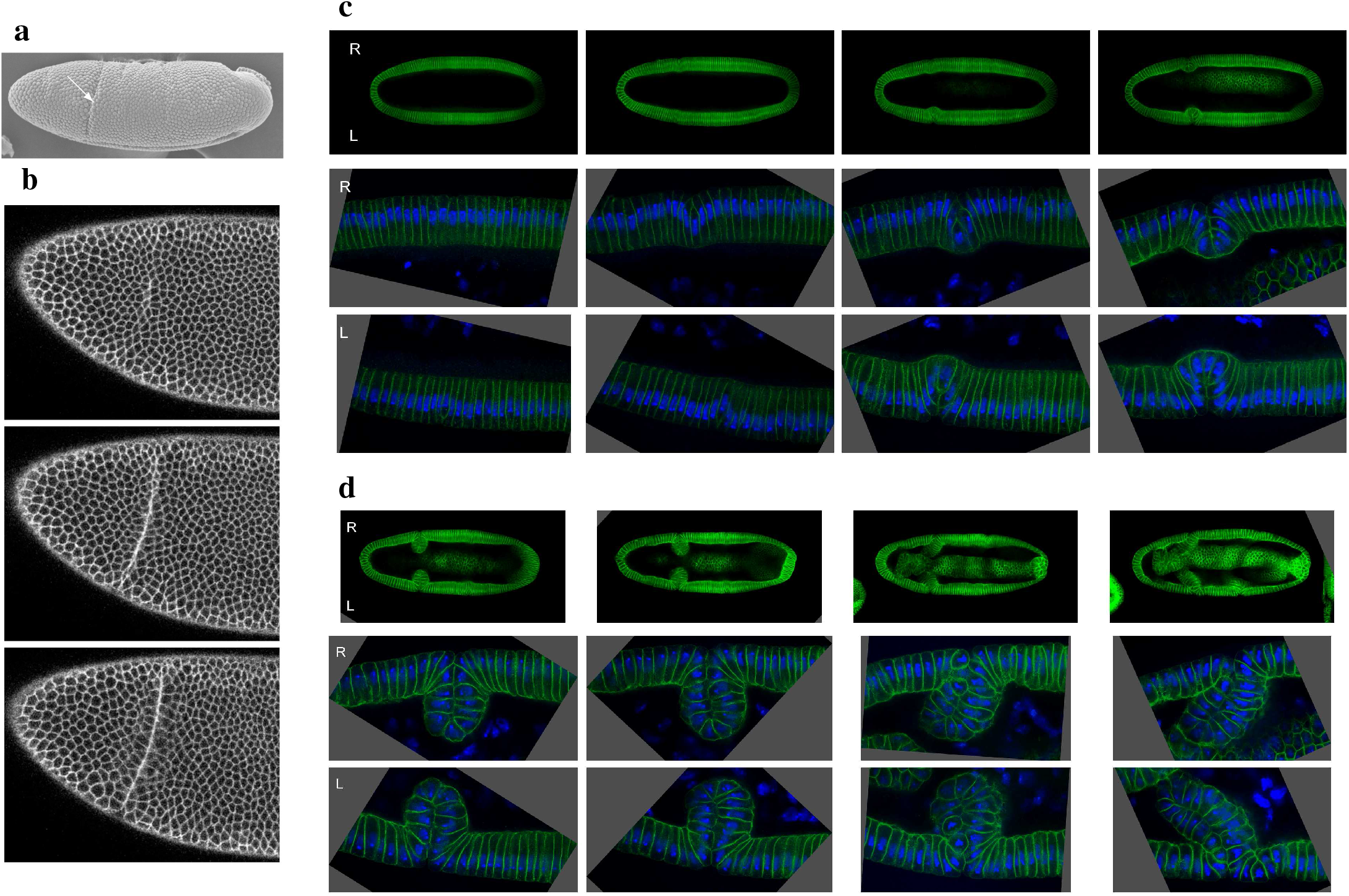
Initiation and progression stages of CFF. (*a*) Electron micrograph of a gastrulating *Drosophila* embryo. The arrow indicates the position of CF. (*b*) Confocal time-lapse images of a Spider-GFP embryo show development of the CF cleft. (*c,d*) A sequence of confocal images of fixed embryos during the (*c*) initiation stage and (*d*) progression stage of CFF. The cell membranes (Nrt, green) and nuclei (Hoechst, blue). During the progression stage consecutive pairs of cells enter the furrow in a semi-periodic manner. All embryo images from Spencer *et al*. (4).

While formation of VF and PMG results in internalization of mesodermal and endodermal precursor cells (5), respectively, CF is not associated with any specific cell fate and does not generate a new germ layer. Moreover, CF is a transitional structure: it has a unique feature in that it first folds, moving cells from the embryo surface into the yolk sac, and then unfolds after the completion of germ band extension (6). Since the transient CF structure forms at the juxtaposition of the head and trunk patterning systems (7), it is likely that its role is to mechanically decouple the head and trunk morphogenetic zones, allowing their robust independent development without mutual interference. Since mechanical forces and feedback strongly affect morphogenesis (1, 8−19), it is crucial to understand forces that govern the CF invagination process.

Most of the previous investigations of the invagination mechanics in the *Drosophila* embryo were focused on VFF, which has been studied in great detail (12, 13, 18−27). It has been shown that, in addition to stress-correlated apical constrictions (12, 13, 27−29), the invagination also involves other mechanical contributions, such as basal expansion and apicobasal shortening of invaginating cells (23), anteroposterior tension along the curved ventral region (18, 19, 27), and possibly a push from the lateral tissue (24, 25). These features were investigated both *in vivo* and via numerical simulations, elucidating how local (23, 24) and long-range (18, 19) mechanical forces shape the embryo, how mechanical cell activity is coordinated through tensile-stress feedback, and how this coordination enhances the robustness of the invagination process (12, 13, 27). However, there are no corresponding investigations of the mechanics of CF formation (CFF).

Unlike VF and PMG formation, CFF is not driven by the coordinated apical constrictions that result in buckling of the invagination region inwards. Instead, mechanical cell activities take place along a single cleft line (Fig. 1, *a* and *b*), where cells enter the furrow in a sequential process (4, 7). As demonstrated by Spencer *et al*. (4), the furrow formation involves subtle cell-shape changes (Fig. 1, *c* and *d*), which are accompanied by myosin II disassembly and F-actin restructuring.

In a recent study it was shown that the fluctuations of the CF cleft line are reduced via mechanical stress (30). However, no detailed mechanical model that realistically takes into account the observed geometry of the furrow-forming cells has been proposed so far. Our present paper fills this gap: we develop an advanced cross-sectional vertex/membrane tension model, which, in addition to the usual balance of pressure and membrane tension forces, also incorporates capillary stresses associated with the membrane tension and curvature.

## MATERIALS AND METHODS

Parental *Drosophila* and their embryos carrying the Spider-GFP transgene were maintained and visualized at 22.5°C (31). Spider-GFP is localized to the cell membranes and thus outlines cell shapes. Embryos were selected at phase 2E in halocarbon oil 27 (Sigma) and prepared for microscopy as described (4, 12, 28, 32, 33). Halocarbon oil was removed, and embryos were mounted by gluing the embryos in chambers that avoided any compression of the developing embryo (4, 12, 13). Time-lapse imaging used a 40x oil objective (NA 1.30) at 15 s intervals. Imaging was conducted using a Nikon Ti-E microscope with A1^+^ confocal.

All images of fixed and stained embryos were data obtained from Spencer *et al*. (4), where details about the imaging methods used are provided. Electron micrograph in Fig. 1 *a*, time-lapse images in Fig. 1 *b*, and Movie S1 in the Supporting Material are also from Spencer *et al*. (4).

## RESULTS AND DISCUSSION

### Key experimental observations

As illustrated in Fig. 1, the CFF process can be divided into two geometrically and mechanically distinct stages: the initiation and progression. The initiation stage (Fig. 1 *c*) corresponds to phase 1 and phase 2E in the nomenclature used in Spencer *et al*. (4). At the beginning of phase 1, initiator cells are undergoing apical constriction and apico-basal shortening (4), which cause the apices of the adjacent cells to meet. Next, these adjacent cells also undergo apico-basal shortening. During phase 1 the CF has not yet started to invaginate into the yolk sac (4). The initiator cells and adjacent cells bulge into the underlying yolk sac during phase 2E (the early part of phase 2), after undergoing basal expansion that initiates the invagination (4). These phases are defined by geometric analyses of cell shape and cellular movements occurring during morphogenesis of the blastoderm epithelium (4).

In the progression stage (Fig. 1 *d*), which is the focus of our investigation, several cells have already invaginated, forming a structurally developed furrow. The progressive stage corresponds to phase 2L, the latter part of the CF invagination phase (4). At this stage, the subsequent cells are added to the furrow in a quasi-periodic iterative process. At the beginning and end of the cycle the invagination region has a similar geometry, but the cells are shifted to the next position in the queue because one cell pair has been added to the furrow.

At the beginning of the progression cycle, the approaching cells assume a wedge-like shape as a result of apical expansion and basal shortening. As the apical ends of the approaching cells meet, the apical membranes roll over each other. The plasma membranes of the two apical sides are now closely juxtaposed as the cells enter the furrow. Concurrently, the pair of cells that met in the previous cycle move further into the yolk sac as their wedge shape transitions into a columnar shape perpendicular to its original orientation. The pair of cells then leaves the active zone of cell shape changes and descends into the furrow.

A single cycle of the progression stage is shown in Fig. Both the initiation phase and the progression phase over multiple cycles are presented in Movies S1 and S2 in the Supporting Material.

A detailed inspection of the magnified image of the active zone where the cell-shape changes occur shows that some lateral cell membranes are curved (Fig. 3). The membrane curvature is indicative of a pressure difference across the membrane (similar to the pressure difference across the membrane of an inflated balloon). The cell membrane curvatures point toward the center of the invagination zone (i.e., the region of wedge-shape cells that undergo rapid shape changes). This geometry indicates that in the center of the invagination zone there is a low pressure region, and it is surrounded by higher pressure domains of the incoming cells and the already invaginated cells (Fig. 3).

**Figure 2:**
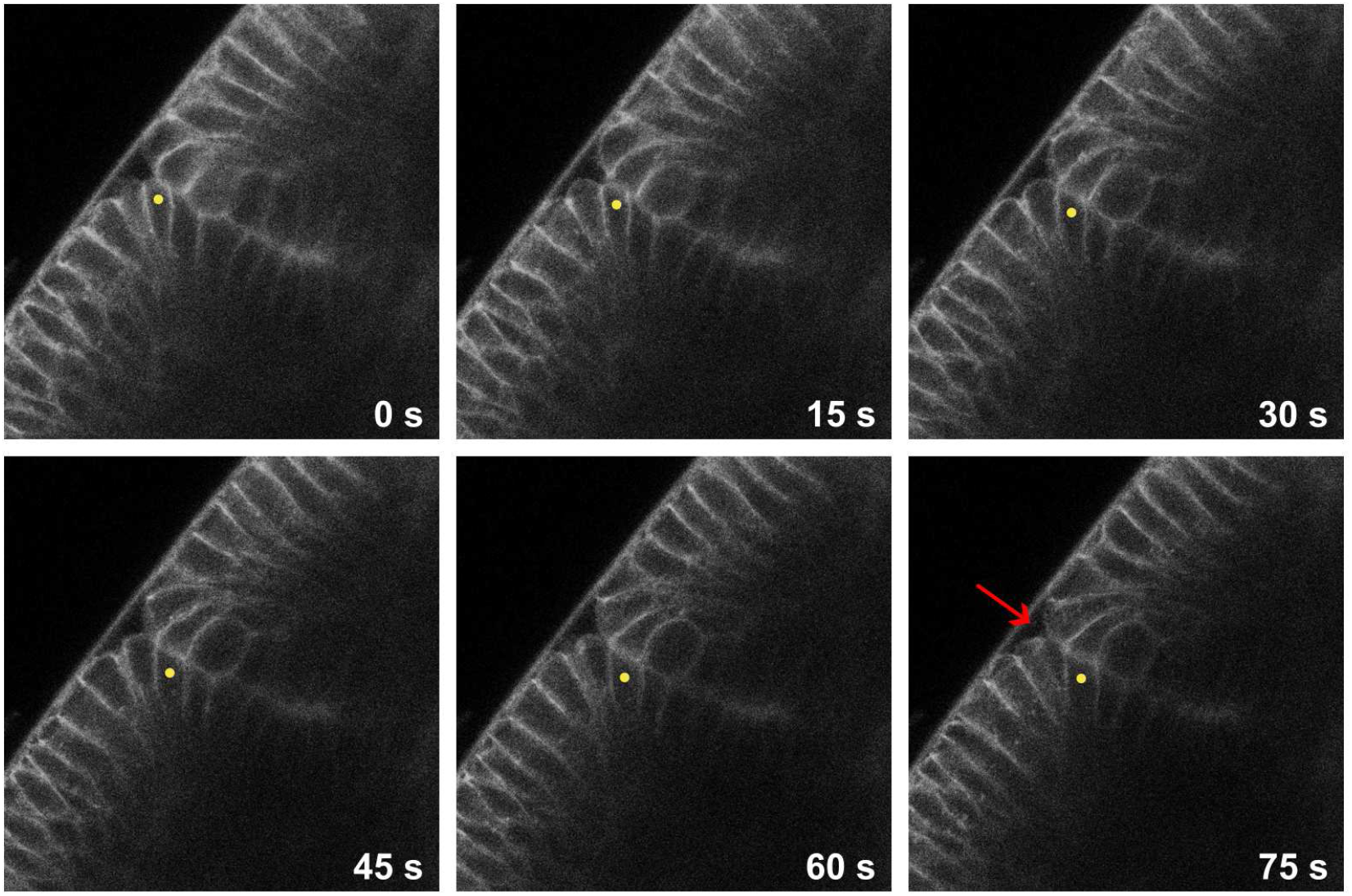
Time-lapse images showing one cycle of the progression stage of CFF. Apical surfaces of the anterior and posterior edge cells meet and the cells enter the furrow (see nomenclature used in Spencer *et al*. 2015). The cells move deeper into the furrow before becoming invaginated cells that are oriented perpendicular to the blastoderm. Images show cell shape changes and timing of events during the transition to invaginated cells. Images are contrast-enhanced. The anterior invaginating cell is marked by a yellow dot, and the Plateau-border-like perivitelline fluid region at the CF cleft is indicated by the red arrow in the last panel.

**Figure 3:**
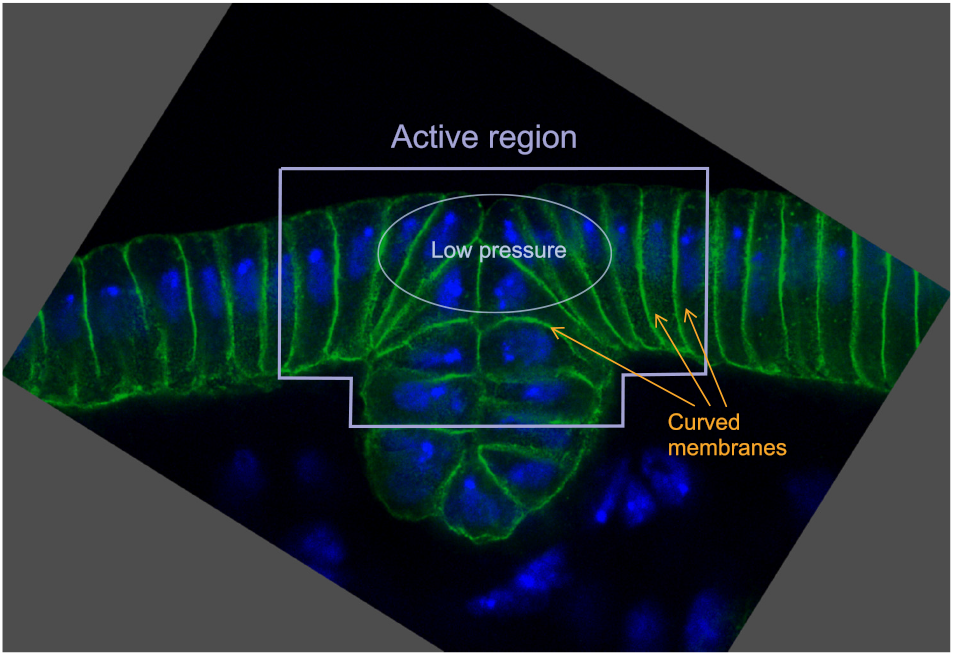
The geometry of the active region. Cell membrane curvature implies the existence of a low pressure region near the CF cleft. Image from Spencer *et al*. (4).

The observed geometrical features of CF provide important clues that can be used to deduce mechanical forces underlying the invagination process. In the next section we develop an advanced vertex model to study the invagination mechanics and the forces involved.

### Advanced cross-sectional model of CFF

#### The geometry of the vertex—spring system

To describe the mechanics of CFF, we developed an enhanced 2D cross-sectional vertex model in which cell membranes are represented by multiple spring segments. As in the standard vertex model (23, 34−36), each cell in the epithelial layer is defined by four main vertices representing the apical− lateral and basal−lateral membrane meeting points. These four vertices form the borders of the apical, basal, and two lateral sides of the cells (the left and middle panels of Fig. 4 *a*). However, unlike the standard approach where each of the apical, lateral, and basal vertex pairs is connected via a single spring representing the cell membrane, we use additional nodes to account for the membrane curvature (right panel of Fig. 4 *a*). If there is a pressure jump across a membrane defined by these multiple spring segments, the membrane bends (the right inset in Fig. 4 *a*), similar to the curved membranes observed *in vivo* (Fig. 3). In this way, the cell geometry can be represented more accurately and the forces driving the invagination determined more reliably. (A similar idea was introduced in (37) to describe packings of deformable particles).

**Figure 4:**
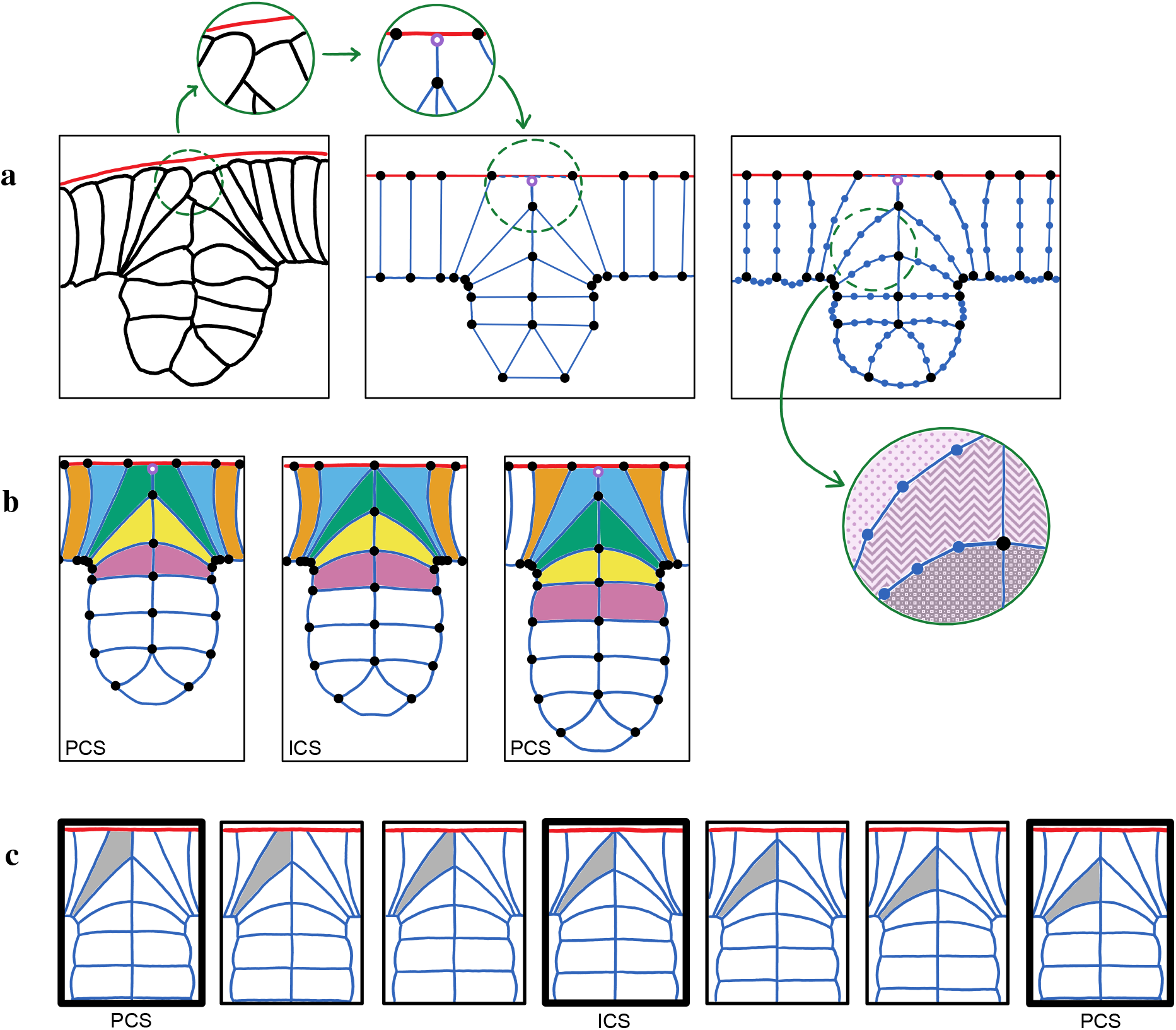
A schematic showing key details of our computational model of the progression phase of CFF. (*a*) The enhanced vertex model. Left panel: outline of cells in the modeled invagination zone. Cell membranes are depicted by black lines and the vitelline membrane by the red line. Middle: cells are described using nodes (black circles) and springs (blue lines). The nodes that represent apical membranes of non-invaginated cells move on a fixed line representing the vitelline membrane (red). The insets show how the perivitelline-fluid region between the vitelline membrane and the innermost anterior and posterior cells at the CF cleft is modeled as a single pulley point (open violet circle). Right: springs representing cell membranes are divided into several segments connected at intermediate nodes (blue points) to account for the membrane curvature associated with the pressure jump from cell to cell (in the inset darker fill color represents higher pressure). (*b*) A set of control states characterizing one period of the progression phase of CFF. The panels show the primary control state at the beginning of the cycle (left); the intermediate control state when a new pair of cells meets (middle); and the primary control state at the end of the cycle (right). The last control state is identical to the initial control state of the following cycle. (*c*) Interpolation of the system parameters between consecutive control states (indicated by the bold frames) followed by equilibration of the node positions generates the system evolution. One of the cells is highlighted to visualize its position and shape changes.

The invaginating part of the epithelial layer as well as the cells approaching from the anterior and posterior directions are modeled explicitly. The remaining epithelial tissue is incorporated implicitly via forces acting on the boundary nodes of the explicitly simulated domain.

The epithelial cell layer that forms the embryo during the CFF process is pressed towards the vitelline membrane by the internal yolk-sac pressure. As a result, apical ends of non-invaginated cells slide along the membrane during CFF. To reflect this behavior in our model, the apical nodes of approaching cells are constrained to move only along the horizontal line representing the membrane position (Fig. 4 *a*).

When two approaching cells meet *in vivo*, their apical membranes roll over each other, with a small amount of low-pressure perivitelline fluid collected in an approximately triangular region between the vitelline membrane and cell membranes at the invagination cleft. This behavior is seen in Figs. 2 and 3 and in the left panel of Fig. 4 *a*. This structure is analogous to the Plateau border in a soap-film foam, i.e., the zone where the interfaces of three adjacent bubbles meet (38).

Here we approximate this Plateau-border-like membrane configuration by a single meeting point (see the left and middle insets in Fig. 4 *a*), similar to a representation of the Plateau border in dry foam simulations (39). The portions of the membrane before and after the meeting point are modeled using a single continuous spring (analogous to an elastic string moving over a pulley). After passing the meeting point, the apical membranes of the anterior and posterior cells of a meeting pair are closely opposed because they are pressed together by the yolk sac pressure. These joint membranes are modeled by a double spring with a common vertex. The fully invaginated cells also have common apical vertices, with a double spring between them. To simplify the representation of the invagination region we have assumed that the system is symmetric with respect to the plane passing through the furrow and normal to the anterior-posterior axis. Thus, the double springs are characterized by a spring constant that is twice the value of an individual spring constant.

#### Interaction potentials

Similar to the approach used in standard vertex models, the dynamics of our enhanced model is governed by linear-spring and surface-compressibility potentials,

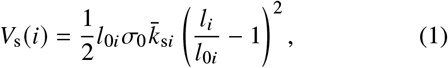

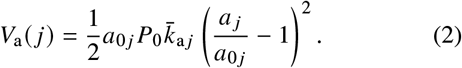

Here *l*_*i*_ and *l*_0*i*_ are the actual and rest lengths of spring *i, a* _*j*_ and *a*_0 *j*_ are the actual and rest areas of cell *j*, and 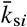 and 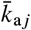 are the dimensionless spring and area elasticity constants. The magnitude of the spring potential, Eq. 1, is set by the membrane-tension scale *σ*_0_, and the magnitude of the area potential, Eq. 2, is set by the pressure scale *P*_0_.

The tensions *σ*_*i*_ acting along cell membranes and the pressures *P* _*j*_ inside individual cells are calculated from the expressions

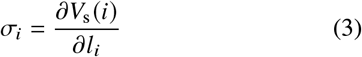

and

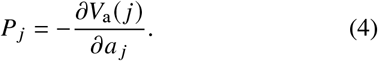

For the harmonic spring and area potentials, Eqs. 1, and 2, we have

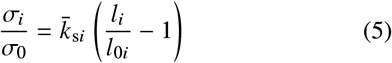

and

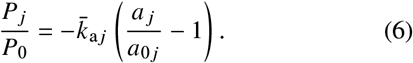

The membrane-tension and pressure scales define the length scale *L*_0_ = *σ*_0_/*P*_0_, which is used to nondimensionalize the cell geometry. In our simulated system the length of the apical membrane before the onset of the invagination process and far away from the invagination region is 1.5*L*_0_.

The force exerted on the explicitly simulated CF domain by the adjacent anterior and posterior epithelial tissue is modeled using the boundary force potential

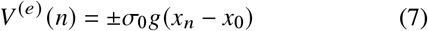

acting only on the cells on the border of the simulation domain. Here *g* is the force strength magnitude, *x*_*n*_ is the position of a border node *n* along the anteroposterior axis, and *x*_0_ is a reference position. The plus (minus) sign applies to the cells on the left (right) boundary of the simulation domain. The boundary force mimics the elastic interaction between the explicitly and implicitly simulated parts of the epithelial tissue, with *g* > 0 corresponding to tension in the epithelial layer.

The evolution of the vertex-spring system representing the CF region is generated by the time variation of the linear-spring parameters 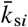 and *l*_0*i*_, area parameters 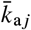 and *a*_0 *j*_, and tissue tension parameter *g*. It is assumed that the evolution is quasistatic, i.e., the system evolves through a succession of mechanical-equilibrium states. Accordingly, at each simulation step the potential of the system

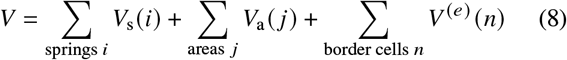

with the current values of the system parameters is minimized to achieve mechanical equilibrium.

The total force at a given node *m* is described by the expression

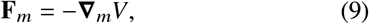

where ∇_*m*_ denotes the gradient with respect to the position of the node. The total forces at the nodes have spring tension, area pressure, and tissue tension contributions. Since we consider a quasistatic evolution, these contributions balance at each simulation step, i.e., **F**_*m*_ = 0 for all *m*.

To minimize the number of independent parameters, the spring and area elasticity constants 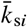 and 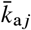 are kept unchanged throughout the entire invagination process, and the tissue tension parameter *g* is set to zero, unless explicitly stated otherwise.

#### Control states

To simulate the progression phase of the invagination process, we first construct a set of control states that mimic the observed cell geometry at key time points of the invagination process. The parameters of the intermediate states of the evolving system are then determined using interpolation.

According to our experimental observations, during the progression stage the invagination occurs in a quasi-periodic manner. In a given invagination cycle one new pair of cells enters the furrow. Each invagination cycle occurs in two distinct phases. In the first phase the cells of a new pair approach each other and at the end of this phase the apical cell membranes meet. In the second phase they roll over each other (Figs. 1 *d*, 2 and movies S1, S2). To mimic this process we use two kinds of control states: (i) the primary control state (PCS) at the beginning of each invagination cycle (left panel of Fig. 4 *b*) and (ii) the intermediate control state (ICS) in the middle of the cycle at the time point when the two approaching cells first meet (middle panel of Fig. 4 *b*). The third (final) control state of each cycle is the same as the PCS of the next cycle (right panel of Fig. 4 *b*).

To establish a control state, we vary the rest lengths *l*_0*i*_ of each spring and rest areas *a*_0 *j*_ of each cell until the system in mechanical equilibrium approximately matches the geometry seen in images of the invaginating region *in vivo*. We semi-quantitatively match such details as the cell shape (columnar vs. wedge-like), cell aspect ratio, and the curvature of lateral membranes. We also match the cell orientation and position with respect to other cells. Since all anterior and posterior cells away from the invagination region are approximately identical, we maintain fixed values of parameters *l*_0*i*_ and *a*_0 *j*_ for all approaching cells that are far from the active domain. Similarly, all columnar cells that have already invaginated have the same values of these parameters. An invagination cycle is complete when the number of invaginated columnar cell pairs increases by one.

#### System evolution

In this section we describe a basic version of our algorithm for evolving the system over multiple cycles of CFF progression phase. In this algorithm, the evolution of the spring-vertex system is achieved by interpolation of all spring-length and cell-area parameters *l*_0*i*_ and *a*_0 *j*_ between their values at the control states. Some necessary modifications of the algorithm for investigations of perturbed systems are described in section “The effect of tissue tension on the invagination process”.

To characterize the progression of the invagination process, we introduce the progression variable *s*. The integer values of this variable,

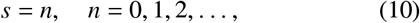

describe the primary control states at the beginning of each invagination cycle, and the half-integer values,

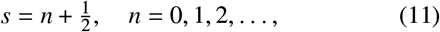

are associated with the intermediate control states when a new pair of approaching cells meets in the middle of the cycle. For uniform progression in time, the variable *s* can be interpreted as the normalized elapsed time *s* = *t*/*T*, where *t* is the time measured from the beginning of the progression phase of CFF and *T* is the duration of a single cycle corresponding to the invagination of a consecutive pair of cells. In the analysis presented below we assume that the progression phase starts when the first pair of columnar cells has invaginated, and we set *s* = 0 at this time instant.

In our interpolation-based evolution procedure we distinguish two subintervals for each invagination cycle—before and after the invaginating cells meet (Fig. 4 *c*), similar to the behavior observed *in vivo*. In the first subinterval of *n*^th^ invagination cycle, 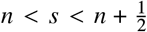, we interpolate between the rest spring lengths 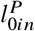 and rest cell areas 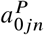 of the primary control state *n* at the beginning of the cycle and the corresponding parameters 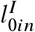 and 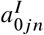 of the intermediate control state *n* in the middle of the cycle,

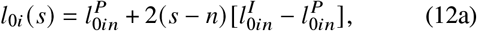

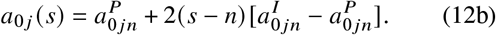

Here *l*_0*i*_ (*s*) and *a*_0 *j*_ (*s*) are the rest length of spring *i* and the rest area of cell *j* at normalized time *s*. In the second sub-interval period, 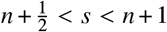, we interpolate between the intermediate control state *n* and the primary control state *n* + 1 at the end of the cycle,

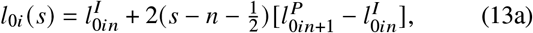

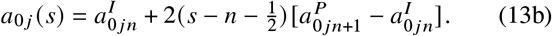

When switching between the interpolation formulas 12 and 13, the topology of the system is restructured by moving the apical membranes of the meeting cells over the Plateau-border pulley point and by creating a new common basal membrane vertex. This procedure corresponds to the formation of a double apical membrane when the cell membranes roll over each other.

In our simulations, each invagination period is divided into small subintervals defined by an increment of the variable *s*. After each interpolation step the system is fully equilibrated by minimizing the potential energy Eq. 8 using the conjugate gradient method.

### Simulation results

Figure 5 and Movie S3 show results of our benchmark simulation of the CF invagination process under conditions where the tension of the tissue that surrounds the explicitly simulated invagination region vanishes, *g* = 0. To reduce the number of free parameters, the dimensionless elastic constants for all springs and areas are set to 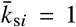 and 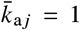. The control states are obtained by adjusting the rest lengths and rest areas *l*_0*i*_ and *a*_0 *j*_ of all cells to approximately match the observed geometry of the CF invagination region. The intermediate states between the control states are determined by interpolation, Eqs. 12 and 13.

**Figure 5:**
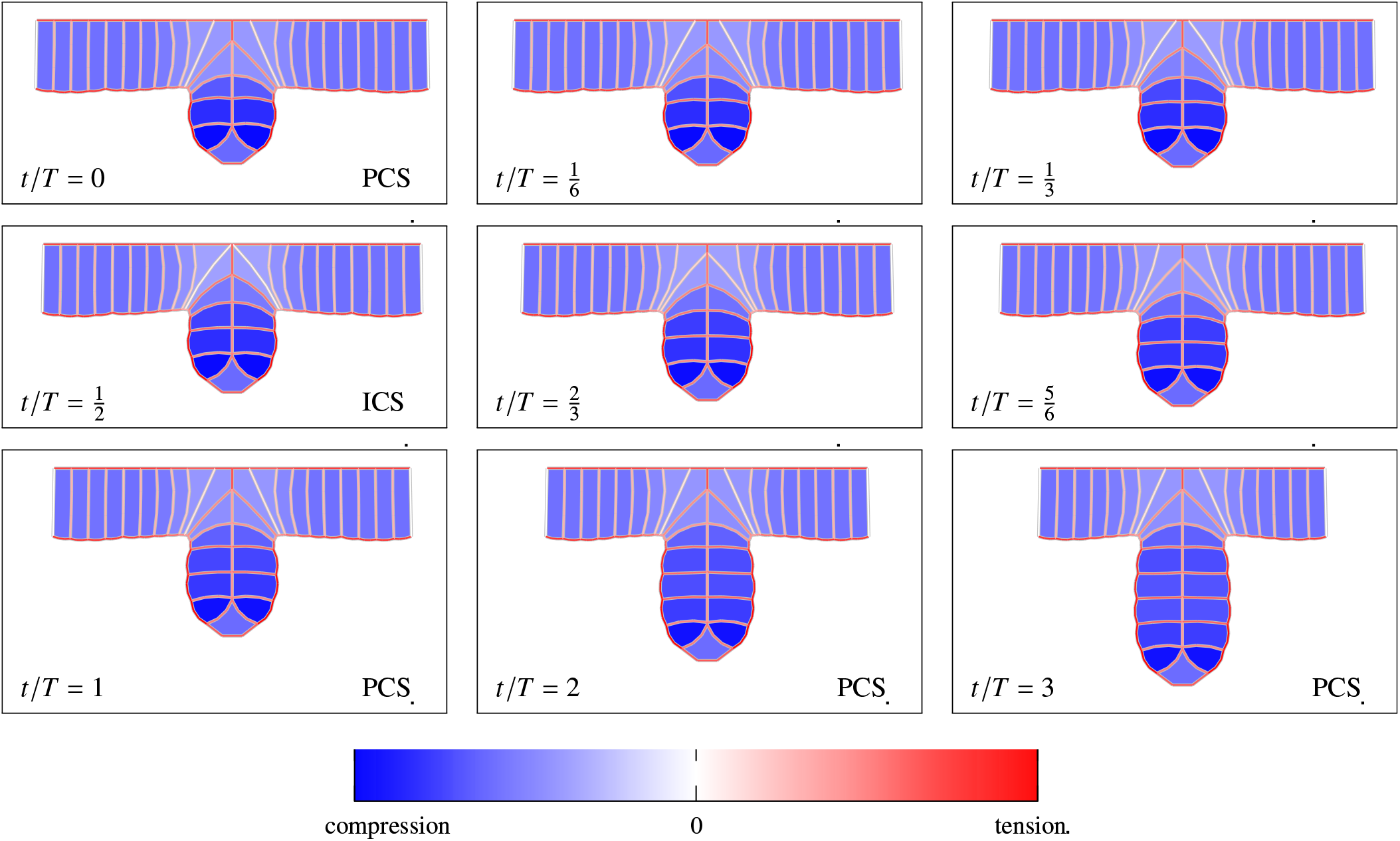
Benchmark (*g* = 0) simulation of the progression stage. The first seven frames depict a full invagination cycle, with two interpolated states shown between the control states. The next two panels present the primary control states of the two following cycles. Normalized time, as labeled. The primary and intermediate control states are marked PCS and ICS, respectively. The color represents the magnitude of spring tension and area compression, as indicated by the color bar.

In the simulation frames shown in Fig. 5 the spring tensions and cell pressures are indicated using a color scale where blue denotes compression and red tension. The variation of the cell pressure *P* and the changes of the length *l* ^(*α*)^ and tension *σ*^(*α*)^ of the apical (*α* = a), lateral (*α* = l), and basal (*α* = b) membranes are plotted in Figs. 6 and 7. The pressure, membrane length, and membrane tension are normalized by their corresponding values *P*_*∞*_, 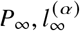, and 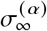 at the boundary of the explicitly simulated region.

**Figure 6:**
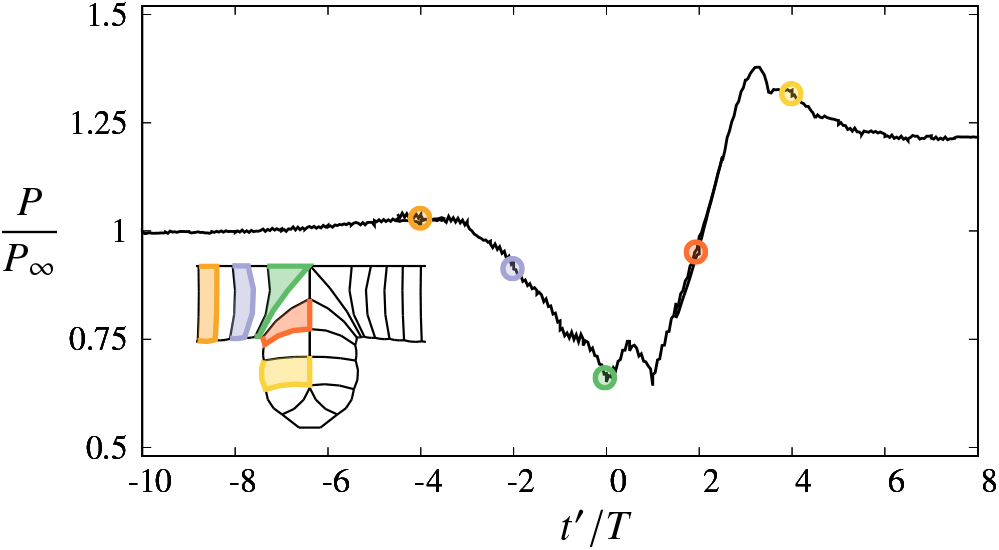
The time variation of the pressure *P* of a cell under-going CF invagination. The results for the system depicted in Fig. 5. The inset shows the cell position at the points marked by the circles of the same color. The pressure is normalized by the value *P*_*∞*_ at the boundary of the explicitly simulated region. The simulations were carried out over seven invagination cycles, and the graph was obtained by combining the results for three cells with different initial positions. The shifted time *t*^′^ is set to zero when the tracked cell reaches the CF cleft. The simulation shows that there is a significant pressure decrease in the active invagination zone.

**Figure 7:**
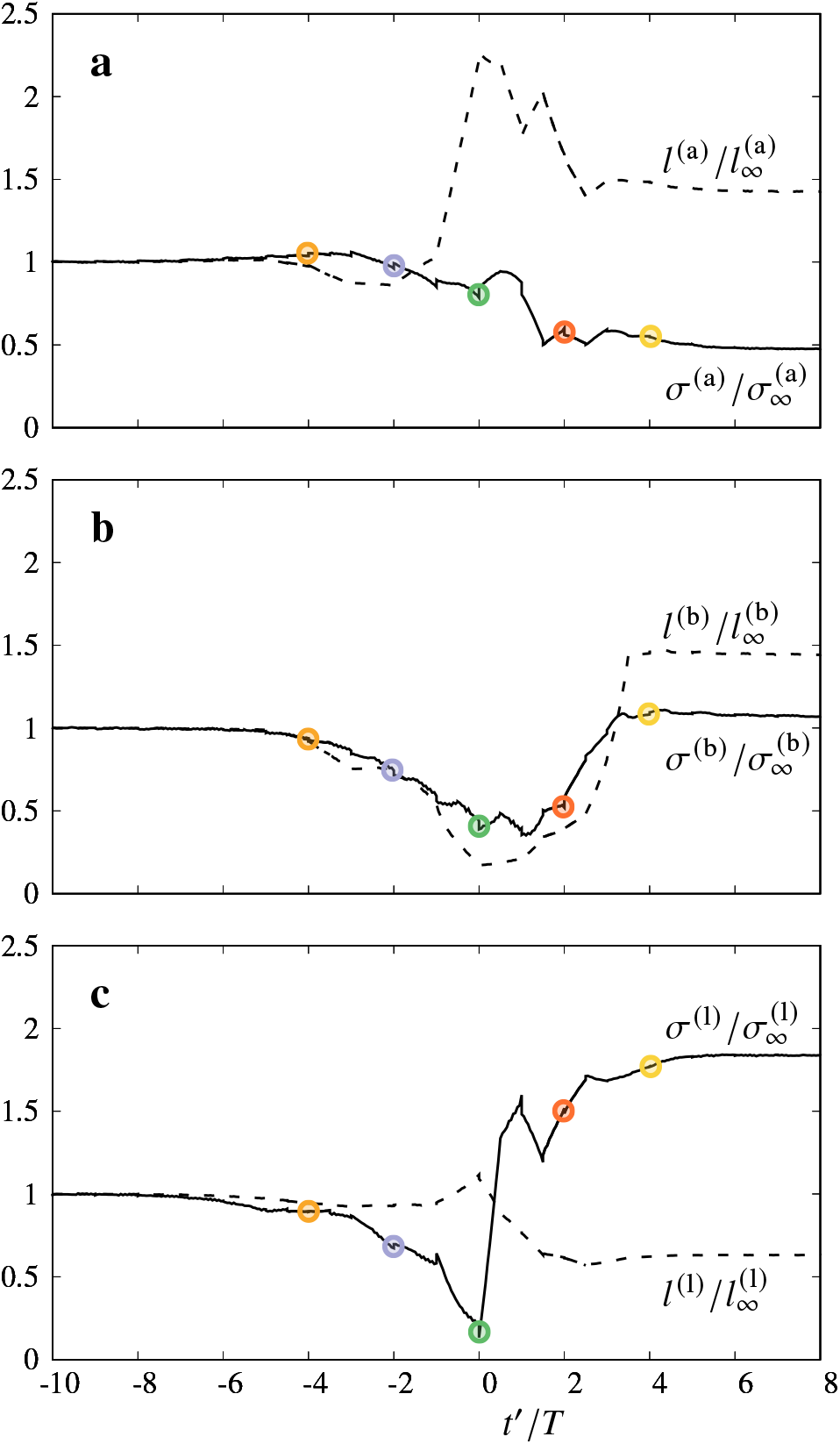
The time variation of the tension *σ* and length *l* of the (*a*) apical, (*b*) basal, and (*c*) lateral membranes of a cell undergoing CF invagination. The results for the system depicted in Fig. 5. The color circles correspond to cell positions indicated in the inset of Fig. 6. The tension and length are normalized by their values at the boundary of the explicitly simulated region. The plots combine the results for three cells with different initial positions. The shifted time *t*^′^ is the same as in Fig. 6. The length variation of the basal membrane closely follows the pressure (Fig. 6) and basal tension variation; this behavior does not occur for the apical and lateral membranes.

#### Geometrical transformations of invaginating cells during CFF

The active cells in the simulated CF depicted in Fig. 5 undergo a similar sequence of geometrical transformations (changes of cell shapes and relative cell positions) as the ones observed *in vivo* (Figs. 1 *c* and 2 and Movies S1, S2). During several progression-phase cycles, the cells entering the active zone transform from the initial columnar shape to the wedge-like shape and then revert to the columnar shape (but more compact and rotated by 90°) when they are fully invaginated.

The membrane length changes (Fig. 7) are consistent with the above transformations of the cell geometry. These length changes include apical expansion followed by apical shortening, basal shortening followed by basal expansion, and elongation and shortening of lateral membranes. During apical expansion the approaching wedge-shaped cells meet at the CF cleft; then they dive into the furrow, rotating by 90°.

The membrane curvatures in our simulation (Fig. 5) reflect those observed in our embryo images (Figs. 1 *d* and 3). Consistent with the curvature orientation seen *in vivo* (i.e. the curvature pointing away from the active domain), there is a low pressure region in the center of the invagination domain, as plotted in Fig. 6 and indicated by the light blue color of cell areas in Fig. 5. To understand forces responsible for the above cell-shape dynamics, we next analyze the cell-pressure changes and behavior of membrane tension.

#### An analysis of the invagination mechanics

The tension of a given membrane results from the balance of the cell-pressure forces and membrane tension forces of the adjacent cells. Thus, the tension of a membrane depends not only on its own mechanical activity but also on the collective behavior of the surrounding cells.

Due to the elongated columnar shape of the cells far away from the active region, the tension of the shorter apical and basal membranes is significantly larger than the tension of the more elongated lateral membranes, as revealed by the deeper color saturation of the corresponding lines in Fig. 5. This behavior stems from the fact that apical and basal membrane tension must balance the pressure force acting on the longer lateral membranes, and this force is proportional to the membrane length (Fig. 8 *a*). For the simulated system, our model yields 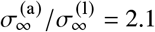 and 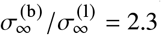.

**Figure 8:**
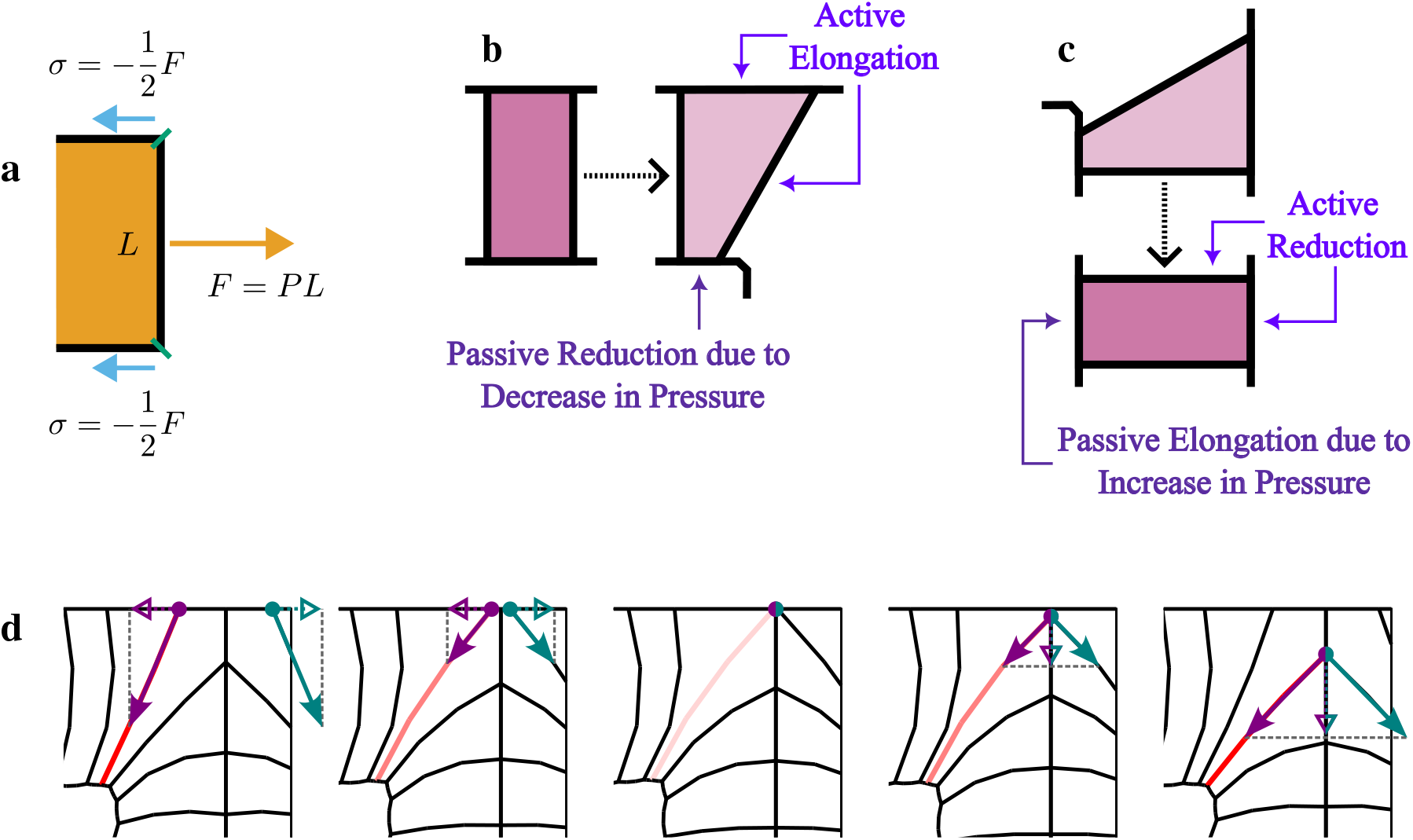
Mechanical forces involved in CFF. (*a*) Tension *σ* of the apical and basal membranes balances the pressure *P* acting on the more elongated lateral membranes; thus the apical and basal tension is larger than the lateral tension. (*b*) Active expansion of the apical and lateral membranes causes passive shortening of the basal membrane in response to the decreased pressure. (*c*) In the reverse process active reduction of the apical and lateral membranes causes passive expansion of the basal membrane in response to the increased pressure. The cell thus returns to the columnar shape. (*d*) Before the cells meet at the furrow cleft, the longitudinal component of the lateral tension force hinders CFF progression (left panels); after the cells meet (middle panel), the transverse component (normal to the embryo surface) promotes the invagination, pulling the cells into the furrow (right panels). The arrow length and the color saturation of the pink line representing the lateral membrane of one of the meeting cells describe the membrane tension variation during the cell encounter.

When a cell enters the invagination region, its apical membrane and the lateral membrane facing the furrow expand, and the basal membrane contracts (time −2 < *t*^′^/*T* < 0 in Figs. 6 and 7). Concurrently with these membrane length changes, the cell pressure and the tension of all membranes decrease.

From the pressure and tension behavior we conclude that the transition of a cell from the columnar to the wedge-like shape is driven by the expansion of its apical and lateral membranes rather than by basal contraction: the expansion causes the pressure decrease, and the basal shortening is likely a passive mechanical response to the reduced pressure and the resulting tension decrease (Fig. 8 *b*). If the basal contraction were the main driving mechanism, we would have observed basal tension increase (needed to actively generate basal shortening), and the shortening would result in pressure increase. This, however, is not the observed behavior.

After the cells have met at the CF cleft (the Plateau-border pulley point in our model) the apical and lateral membranes decrease in length (the time interval 0 < *t*^′^/*T* < 2 in Figs. 6 and 7). The apical and lateral contraction generates the increase of cell pressure, which, in turn, produces basal tension increase and basal expansion (Fig. 8 *c*). The observed tension and pressure behavior in this process, inverse to the one in the preceding interval −2 < *t*^′^/*T* < 0, confirms that basal length changes are a response to the pressure and the corresponding basal tension variation. Hence, the initial basal shortening and subsequent basal expansion do not need to involve active forces in the basal membrane, but occur in response to the activity of apical and lateral membranes.

When a lateral membrane approaches the cell meeting point at the CF cleft (*t*^′^/*T* = 0 in Figs. 6 and 7), the membrane tension decreases to a value close to zero. (The membrane, however, does not undergo compression because compression would result in buckling.) After the lateral membranes of the approaching cells meet (i.e., the springs in our model reach the intermediate control state and go over the pulley point) the membrane tension sharply increases. Our simulations demonstrate that the initial tension decrease followed by the increase is needed for a successful invagination.

This behavior is explained in Fig. 8 *d*. Before the approaching cells meet, the tension of the diagonally oriented lateral membranes pulls the cells apart, hindering the invagination process. Thus, the tension must decrease to enable the invagination progression. After the cells meet and their apical membranes start to roll over each other, the lateral tension draws the cells on each side inwards, assisting invagination. The cells on each side of the furrow are connected to the cells directly above and below by adherens junctions near the apices. We note that during this process the apical tension gradually decreases (Fig. 7 *a*) because part of the pulling force exerted by the apical membrane of the following cell is balanced by the diagonally oriented lateral membrane (Fig. 8 *d*).

#### The role of the cell pressure variation

To explain the mechanical role of the pressure decrease when the cells enter the active region, we have generated the intermediate control state for a system with approximately uniform pressure and compared it to the corresponding benchmark system with the low-pressure active region (Fig. 9). To ensure that the approaching cells meet in the hypothetical uniform-pressure case, both a decrease of the rest lengths and an increase of the spring constants were necessary for the apical double springs in the center of the active region. The resulting membrane-tension increase enabled the meeting of the approaching cells, but it also produced a deformation of the invaginated domain (Fig. 9 *b*). The cells in the active region assumed a tilted parallelogramic shape, with apical ends pointing towards the vitteline membrane, unlike the corresponding columnar cells in the benchmark simulations with the pressure decrease in the active region (Fig. 9 *a*).

**Figure 9:**
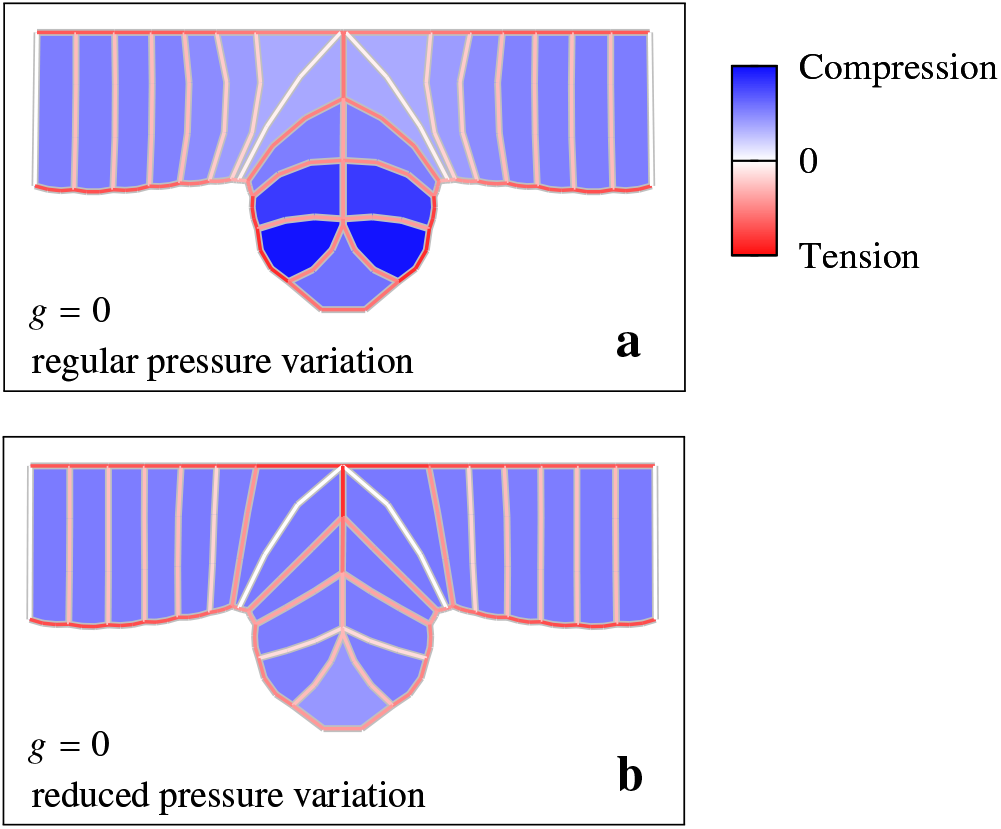
The effect of the pressure distribution. Intermediate control state for a system with *g* = 0 and (*a*) the regular pressure variation and (*b*) no pressure decrease in the invagination region. Without the pressure variation the invaginated cells are tilted towards the center of the active region. The depicted control states correspond to time *t/T* = −1*/*2 on the scale defined in Fig. 5.

The mechanism by which the variable pressure distribution reduces the deformation of invaginating cells includes a combination of the following effects. First, since the low-pressure active cells are more compliant than the surrounding higher pressure columnar cells, it is possible to generate the prescribed shape changes without applying significant forces. Second, the approaching and the already invaginated high-pressure columnar cells form stiff blocks to which a chain of high-tension apical membranes is attached. In conjunction with active cell shape changes (columnar to wedge-like and back to columnar), this apical membrane chain, attached to rigid supports of columnar cells, helps to pull new cells into the furrow without causing deformation of the invaginated region.

### The effect of tissue tension on the invagination process

In this section we ask whether the CF can pull new tissue into the furrow against the tension of the non-invaginated epithelial cells. To address this question, we performed numerical simulations of the system with nonzero tissue tension force *g* in Eq. 7. Since cell meeting at the CF cleft is the critical time point of each invagination cycle, we focus on the effect of tension force *g* on the intermediate control states, i.e., the states with apical cell membranes just touching (Figs. 10 and 11).

**Figure 10:**
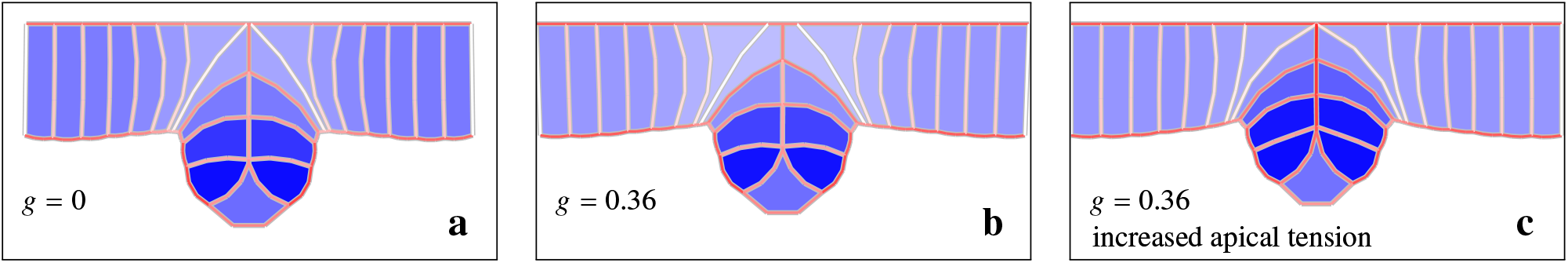
The effect of the anterior and posterior tissue tension *g* on the cell-meeting event. (*a*) A system with apical membra of approaching cells touching before the external tension is applied; (*b*) when the tissue tension force is switched on, the ce separate; however, (*c*) the cells meet again after apical tension in the invagination region has been increased. The simulati frames correspond to the time *t/T* = −1*/*2 on the scale defined in Fig. 5. The applied tissue-tension force *g* is normalized by tension of the lateral membrane of a cell on the border of the explicitly simulated region.

**Figure 11:**
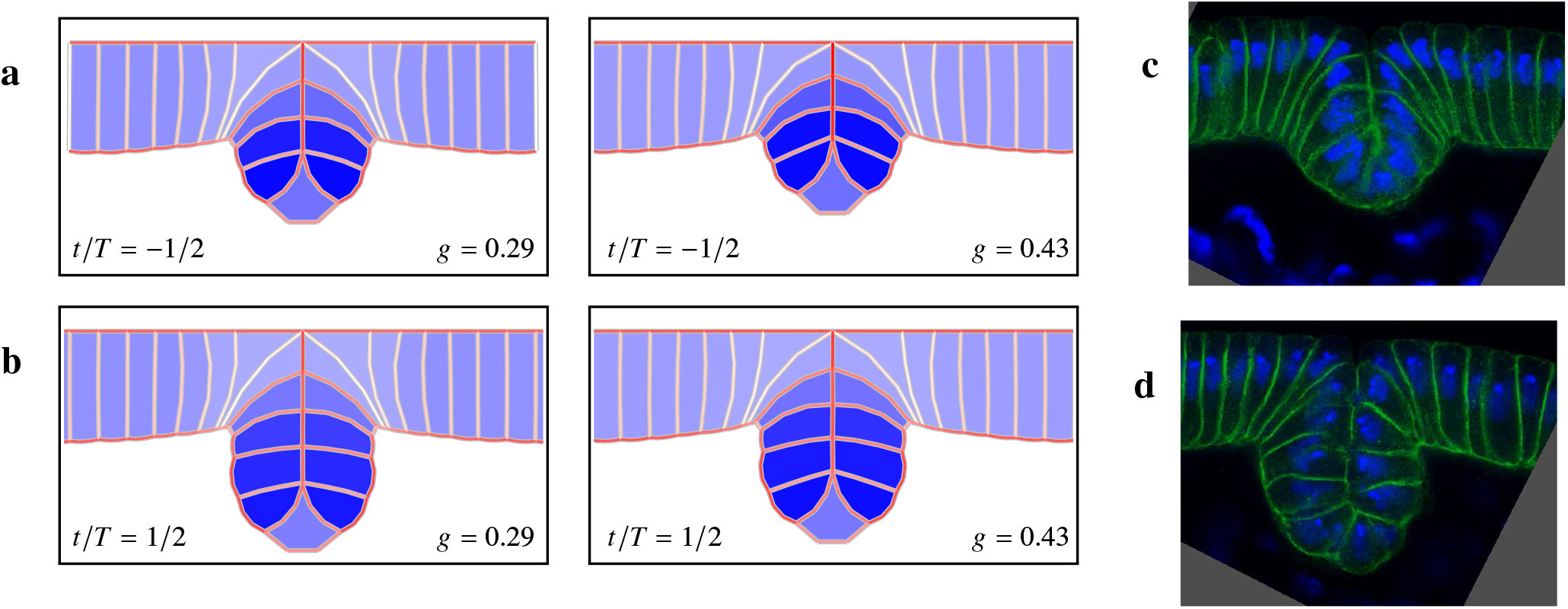
The effect of tissue tension *g* on the geometry of the active and invaginated regions. The simulation frames with apical membranes of approaching cells meeting at the invagination cleft for (*a*) *t/T* = −1*/*2 and (*b*) *t/T* = 1*/*2 and two different values of the normalized tissue-tension force *g* (as labeled). Embryo images (*c*) at the transition from the initiation to the progression stage and (*d*) about one cycle later. The *in vivo* images approximately correspond to the simulation images depicted in panels (*a*) and (*b*). Images (*c*) and (*d*) from Spencer *et al*. (4).

Figure 10 *a* and *b* shows that without an appropriate mechanical activity of the invaginating cells (membrane tension adjustment), the tension in the non-invaginated epithelial layer can impede CFF because the pulling force resulting from tissue tension can prevent approaching cells from meeting. However, after the tension of apical membranes in the active region is increased (by adjusting the spring constant 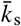 in our simulated system), the cells meet, which restores invagination progression (Fig. 10 *c*). Thus, with a proper active response of cells to applied stresses, the furrow forms in spite of the opposing tensile stress in the layer of the non-invaginated cells.

Figure 11 illustrates how the tissue tension magnitude *g* and the size of the already invaginated domain impact the cell geometry. Two characteristic time points are considered, *t/T* = −1*/*2 and *t/T* = 1*/*2 (on the scale defined in Fig. 5).

The first case (Fig. 11 *a*) corresponds to the transition between the initiation and progression stage; in the second case (Fig. 11 *b*), an additional pair of cells has already entered the furrow. For *t/T* = −1*/*2 the cells entering the furrow undergo significant deformation, with the apical ends pointing towards the surface of the embryo, especially for the larger value of *g*. The deformed configurations are similar to the one seen for the system with reduced pressure variation and *g* = 0 (Fig. 9 *b*); however, the membrane curvature is distinct in these two systems. For *t/T* = 1*/*2, the presence of the additional pair of cells reduces the deformation nearly completely. The invaginated cells have a columnar shape, with only a slight tilt at the stronger tension (larger value of *g*). This stems from the fact that the invaginated high-pressure columnar cells form a stiff base, which helps to pull new cells into a CF groove and then align them normal to the invaginated tissue.

We note that a similar sequence of changes is observed *in vivo*. At the transition between the initiation and progression stages the apical ends of cells entering the furrow point towards the embryo surface and away from the yolk sac, but this tilt is removed after one or two invagination cycles. Based on our simulations and this observed behavior, we hypothesize that there is a nonzero tissue tension in the epithelial layer on both sides of the CF cleft. The role of this tension is discussed in the “Conclusions” section.

## CONCLUSIONS

Mechanical activity in the cell apex is known to be involved in morphogenetic movements of *Drosophila* and most other studied organisms. Apical constriction driven by actomyosin pulses is one of the major morphogenetic activities (40−42). However, a morphological study of CF suggested that cell shape changes involved in this invagination are not driven by apical constriction but by some other mechanism (4). Myosin II is not concentrated at the cell apices of the developing CF but is present in the lateral regions (4, 30). Accordingly, CFF cell shortening involved in formation of the furrow is mediated by actomyosin contraction along the lateral region (30). This and other differences between CFF and other morphogenetic movements provide strong motivation to investigate CF invagination mechanics, using the observed time sequence of cell shapes and furrow morphology.

The CFF process can be divided into two distinct stages: initiation and progression. During the initiation stage (phases 1E, 1L, and 2E in the nomenclature introduced in (4)) the initiator cells and adjacent cells undergo apical constriction and apicobasal shortening, which prepare the region for the upcoming invagination. Subsequently, the epithelial layer starts to bulge into the yolk sack. The progression stage of CFF (phase 2L (4)) is characterized by the rapid and deep invagination of many cells in a semi-periodic process during which subsequent pairs of cells roll over the furrow cleft, undergo a 90° turn, and drop into the interior. The invaginating cells first assume a wedge-like shape, their apical membranes roll over each other, and then the cells return to the columnar shape undergoing apico−basal shortening during this process. In the present paper we examine the mechanics of the progression stage of CFF.

To analyze the narrow invagination of CFF, we developed an advanced 2D cross-sectional vertex model which considers pressure, cell cortex tension forces, and capillary forces associated with cortical tension and curvature. One of the innovations in our advanced 2D vertex model is the use of multiple nodes in representing the lateral and basal cell cortices and plasma membranes. This allows us to more accurately model cell shape changes and capture how cellular pressures vary as the cells enter the furrow and progress towards the embryo interior.

As cells approach the furrow cleft, they change from the columnar to wedge shape by undergoing apical expansion and concomitant basal reduction. Accompanying these membrane length changes, there is a drop in the internal cell pressure, as indicated by the cell curvature observed *in vivo* and reproduced in our numerical model. As the cells rotate and move into the perpendicular fold, their pressure rises close to the previous levels while apical tension drops, basal tension rises slightly, and lateral tension increases about 50% over the original baseline. Once the cell has become fully perpendicularly aligned, the pressure in the cell is 30% higher than before entering the furrow (See Figs. 6 and 7).

The results of our numerical simulations show that the variation in cellular pressure plays a pivotal role in ensuring an orderly progression-phase process. First, the pressure produces mechanical forces that, combined with the membrane-tension forces, help to move subsequent cells into the furrow. Next, the pressure transmits stress information between the apical, lateral, and basal membranes, coordinating their activities.

The direct pressure-force contribution during the initial pressure-decrease phase and the subsequent pressure-increase one has two distinct and complementary functions when a cell moves over the CF cleft and into the furrow. The decreasing pressure in a cell that rolls over the CF cleft softens the cell, thus facilitating the required cell shape changes. In contrast, the gradually increasing pressure during the transition from the wedge-like shape back to the columnar one generates the active force that pulls new cells over the furrow cleft.

Moreover, the higher pressure of the cells that have completed their queue into the deepening furrow provides a solid foundation for the cells that produce the pulling force, thus enhancing the effectiveness of their activity. The foundation of high-pressure cells enables furrow progression even in the presence of opposing forces produced by a nonzero tension of the anterior and posterior tissue.

The above finding points to one of the possible purposes of CFF, i.e., ensuring a proper tension distribution over the embryo. It has recently been shown that tensile mechanical stress feedback plays an important role in VFF (12, 13, 24, 27). Since tensile stress feedback is most effective in an environment where there is already some baseline of tensile stress across the system, we hypothesize that transient CFF may help support mechanical feedback mechanisms vital for other morphogenetic movements by temporarily absorbing some of the embryonic epithelium so that there is adequate tension maintained across the embryo.

Based on the predicted correlations between the cell pressure variation and the apical, lateral, and basal membrane length and tension changes, we propose that wedging of a cell approaching CF cleft is controlled by the apical expansion and the concurrent expansion of the lateral membrane facing the furrow. The expansion, resulting from the release of the membrane tension due to restructuring of the apical and lateral cortex, lowers the pressure in the cell. The lowered pressure produces, in turn, a correlated decrease of basal tension and induces basal reduction. After the cell has entered the furrow, a rearrangement of the apical cortex reduces the size of the apical region, and this reduction is accompanied by a simultaneous contraction along the lateral region. These together produce an increase of the cell pressure, which drives the passive expansion of the basal cortex. The cell thus shortens and returns to the columnar shape.

According to the above analysis, the active elements that govern the shape changes of an invaginating cell are the cell cortices underlying the apical and lateral membranes. This hypothesis is supported by the extensive apical cortical F-actin reorganization initiated by the dynamic expansion of F-actin levels in the apices of invaginated cells during phase 2L (4). The pressure decreases or increases according to the changing size of the apical an lateral membranes, the pressure variation causes the corresponding tension changes in the basal membrane, and the basal membrane contracts or expands in response to the tension changes. The basal membrane thus reacts to the mechanical activity of the apical and lateral membranes, using the stress information transmitted via cell pressure.

Our finding is in line with recent work that suggests that the basal cortex in the *Drosophila* embryonic blastoderm is inactive but responsive to both stress and pressure (42). A passive basal element was also implemented in the 2D Polyakov model of VFF (23). In their model, the only active forces were those involved in apical constriction. Other cell shape changes were suggested to be driven by remodeling of their cortical cytoskeleton in response to the stresses and tensions in the embryo caused by apical constrictions (23). CFF and VFF are distinct morphogenetic movements with a variety of differences; however, they occur around the same time and involved cells are subject to similar physical conditions. Our results, therefore, add to the mounting evidence that the basal cortex of cells during this stage of development is a passive element, which can respond to its mechanical environment.

It was recently shown by another group that genetic control during CFF phase 1E is only partly responsible for the organization and alignment of the initiating cells (30). Using a combination of detailed experiments and a 3D vertex model, they found that approximately 20% of initiator cell alignment results from a response of the tissue to tensile forces that stretch it along the future CF cleft (30). Their results demonstrate the involvement of mechanical tensile stress in CFF.

Our recent work on VFF (27) and work of others (18, 19) suggest that tensile stress oriented along the long axis of the ventral field is essential in the entire VFF process. First, this stress coordinates apical constrictions (12, 13, 27), leading to formation of actomyosin meshwork (18, 28, 43) and generating a narrow band of tissue tension on the curved embryo surface. Second, this tensile stress produces localized inward pressure that causes inward tissue deformation (18, 19). Finally, the tensile stress and the associated inward pressure support mechanical stress feedback that is pivotal in VFF (27).

These results lead us towards the possibility that similar mechanisms also affect CFF. We propose that tensile stress feedback is at play in aligning the initiator cells along tension generated by the newly forming CF groove. In this process, the tensile-stress-generated inward force provides a mechanical feedback signal that initiates the invagination. Later, during the progression phase, cell activities are coordinated along the entire groove by long-range tensile forces and the associated inward force and mechanical feedback.

The above picture is supported by the fact that F-actin is highly enriched in the apices of the cells in the CF cleft and is even seen in phase 1 (4). We expect that F-actin is arranged either into intercellular cables that run along the apices of the invaginating cells or into a less rigid meshwork, both of which are capable of transmitting tensile stress effectively. Confirmation of such F-actin arrangements would suggest that CFF is driven not only by cell shape changes evident in the cross-sectional view of the furrow but is also assisted by tensile stress along the furrow and thus by the resulting mechanical stress feedback and inward force. It is also likely that the lateral actomyosin contractions observed (30) in the initiator cell region also contribute to the tension along the furrow.

Our advanced 2D vertex model of CFF has allowed one of the first in-depth analyses of the cell mechanics during the progression phase. We have demonstrated the importance of cell pressure variations and tissue tension, and provided evidence that the basal cortex is a passive element at this stage in development. The work presented here forms the basis for more detailed analyses of this relatively unexplored morphogenetic process. Further studies of CF invagination are likely to uncover various mechanical stress feedback mechanisms that guide CFF.

## AUTHOR CONTRIBUTIONS

JB and RAN developed the computational model and simulation algorithms. They also analyzed simulation and experimental data. All simulations were performed by RAN. JHT provided the experimental data. MCH, JHT, and JB conceived and developed the project. All authors contributed to data analysis, manuscript writing, and conducted insightful discussions of all aspects of the project.

## MOVIE CAPTIONS

Movie S1. Embryo expressing Spider-GFP in early CF invagination.

Movie S2. Embryo expressing D-Ecad-GFP during CF invagination. (Movie from Spencer *et al*. 2015 (4).)

Movie S3. Benchmark simulation of the progression stage of CFF.

